# Shared mechanisms drive ocular following and motion perception

**DOI:** 10.1101/2023.10.02.560543

**Authors:** Philipp Kreyenmeier, Romesh Kumbhani, J. Anthony Movshon, Miriam Spering

**Affiliations:** Department of Ophthalmology & Visual Sciences, University of British Columbia, Vancouver, BC V5Z 3N9 Canada; Graduate Program in Neuroscience, University of British Columbia, Vancouver, BC V6T 1Z3 Canada; Center for Neural Science, New York University, New York NY 10003, USA; Department of Psychology, New York University, New York NY 10003, USA; Institute for Computing, Information, and Cognitive Systems, University of British Columbia, Vancouver, BC V6T 1Z3 Canada; Djavad Mowafaghian Center for Brain Health, University of British Columbia, Vancouver, BC V6T 1Z3 Canada

**Keywords:** pattern motion, ocular following, eye movements, perception-action

## Abstract

How features of complex visual patterns combine to drive perception and eye movements is not well understood. We simultaneously assessed human observers’ perceptual direction estimates and ocular following responses (OFR) evoked by moving plaids made from two summed gratings with varying contrast ratios. When the gratings were of equal contrast, observers’ eye movements and perceptual reports followed the motion of the plaid pattern. However, when the contrasts were unequal, eye movements and reports during early phases of the OFR were biased toward the direction of the high-contrast grating component; during later phases, both responses more closely followed the plaid pattern direction. The shift from component- to pattern-driven behavior resembles the shift in tuning seen under similar conditions in neuronal responses recorded from monkey MT. Moreover, for some conditions, pattern tracking and perceptual reports were correlated on a trial-by-trial basis. The OFR may therefore provide a precise behavioural read-out of the dynamics of neural motion integration for complex visual patterns.

## Introduction

The primate visual system analyzes motion in two stages. First, orientation- and direction-selective neurons in primary visual cortex (V1) signal the velocities of single components within a pattern. Second, component velocity signals are integrated to compute the true motion of objects and patterns. The perceptual and motor impact of these two stages of motion processing can be conveniently studied using 2-dimensional plaid patterns, created by summing 1-dimensional gratings of different orientation (Adelson and Movshon, 1982). When the grating components of a plaid have equal contrast, human observers perceive the coherent motion veridically. But when one component is higher in contrast, the perceived motion is biased toward it (Stone et al., 1990; Yo and Wilson, 1992; Bowns, 2013). This form of motion integration may reflect the activity of neurons in extrastriate visual area MT (Movshon and Newsome 1996; Rust et al., 2006), a main motion processing hub for perception and oculomotor control (Newsome et al., 1985; Newsome and Paré, 1988; Dürsteler and Wurtz, 1988; Kawano, 1999; Lisberger and Movshon, 1999; Takemura et al., 2007; Ilg and Thier, 2008). Recording experiments in MT revealed “component cells” that are tuned to the direction of single plaid components, and “pattern cells” that integrate component signals and are tuned to the true direction of a plaid (Movshon et al., 1985; Rodman and Albright, 1989; Smith et al., 2005; Rust et al., 2006). When plaid components have different contrasts, the true motion of the pattern is unchanged, but the direction perceived by human observers and tuning of MT pattern cells both shift towards the direction of the higher contrast component (Stone et al., 1990; Kumbhani et al., 2008).

We wondered how the signals that give rise to the perceptual experience of motion are related to those that drive motion-dependent eye movements. The eye movement we chose is the ocular following response (OFR) – a reflexive, short latency eye movement evoked by the onset of large-field visual motion (Miles et al., 1986; Gellman et al., 1990; Masson and Perrinet, 2012). With a latency of ∼85 ms in humans (Gellman et al., 1990), 55 ms in monkeys (Miles et al. 1986), and 50 ms in marmosets (Pattadkal et al., 2023; Yip et al., 2023) the OFR provides a direct, behavioural readout of early visual motion processing (Masson and Perrinet, 2012) and of motion integration over time (Masson and Castet, 2002). We simultaneously recorded human perceptual direction estimates and OFR in response to plaids composed of gratings with varying luminance contrasts and presented for different lengths of time. Similar contrast-dependent pattern sensitivity in eye movements and perceptual reports would be congruent with psychophysical studies concluding that eye movements and perception rely on shared processing of direction and speed signals (Gegenfurtner et al., 2003; Stone and Krauzlis, 2003; for reviews, see Schütz et al., 2011; Spering and Montagnini, 2011). Alternatively, visual signals may be processed differently for eye movements and perception (Spering and Gegenfurtner, 2007; Tavassoli and Ringach, 2010; Spering et al., 2011; Simoncini et al., 2012; Glasser and Tadin, 2014; Lisi and Cavanagh, 2015; for a review, see Spering and Carrasco, 2015).

Our results show that perceptual reports of briefly presented motion and the earliest OFR are biased by the high-contrast component of the plaid. With increasing presentation duration and during later OFR and continuous tracking, perceptual reports and the OFR shift towards the direction of the pattern. During these late phases, perceptual and oculomotor performance are correlated on a trial-by-trial basis. This suggests a tripartite link between visual motion perception, oculomotor control, and the cortical machinery of motion processing.

## Materials and Methods

### Observers

We collected data from 8 observers (mean age 28 y, five female, one author). Observers had normal or corrected-to-normal visual acuity and no history of ophthalmic or neurological disorders. The University of British Columbia Behavioral Research Ethics Board approved all experimental procedures, which were in accordance with the Declaration of Helsinki. Observers gave written informed consent and received a remuneration of 10 CAD/hour.

### Apparatus and stimuli

We conducted the experiment in a dimly-lit laboratory. Observers viewed stimuli binocularly on an 18” (1280×1024 pixels; 85 Hz) gamma-corrected CRT monitor (ViewSonic G90fB; La Brea, CA) placed at a viewing distance of 50 cm at which the screen subtended 38.7° (horizontal) × 30.9° (vertical). A PC running Matlab R2019b (MathWorks Inc., Natick, MA) and Psychophysics (version 3.0.12; Brainard, 1997; Pelli, 1997) and Eyelink toolboxes (Cornelissen et al., 2002) controlled stimulus presentation and data acquisition. We recorded the position of the right eye with an Eyelink 1000 tower mount eyetracker (SR Research, Kanata, ON, Canada) at 1,000 Hz.

We compared perceptual estimates of motion direction with eye movements elicited in response to briefly presented gratings or plaids shown within a suitably vignetted circular aperture subtending 20°. The gratings and plaids had the same mean luminance as the background (61.7 cd/m^2^). Plaids were made by adding two drifting sinewave gratings whose motion directions, always orthogonal to their orientations, differed by 120°. We varied the Michelson contrast of one grating between 2.5 and 40% in octave steps while holding the contrast of the other (high-contrast) grating constant at 40% to create 5 different plaids with contrast ratios 1:16, 1:8, 1:4, 1:2, or 1:1. We also presented a single, high-contrast grating (40% contrast). The computer selected the motion direction of the high-contrast component (red arrow in **Fig. 1A**) randomly on each trial; the other component always moved in a direction 120° counter-clockwise (blue arrow, **Fig. 1A**). The grating spatial frequency was 0.25 c/deg and the drift rate 6 Hz, yielding a speed of 24 °/s for each component grating and 48 °/s for the plaid.

**Figure 1.**
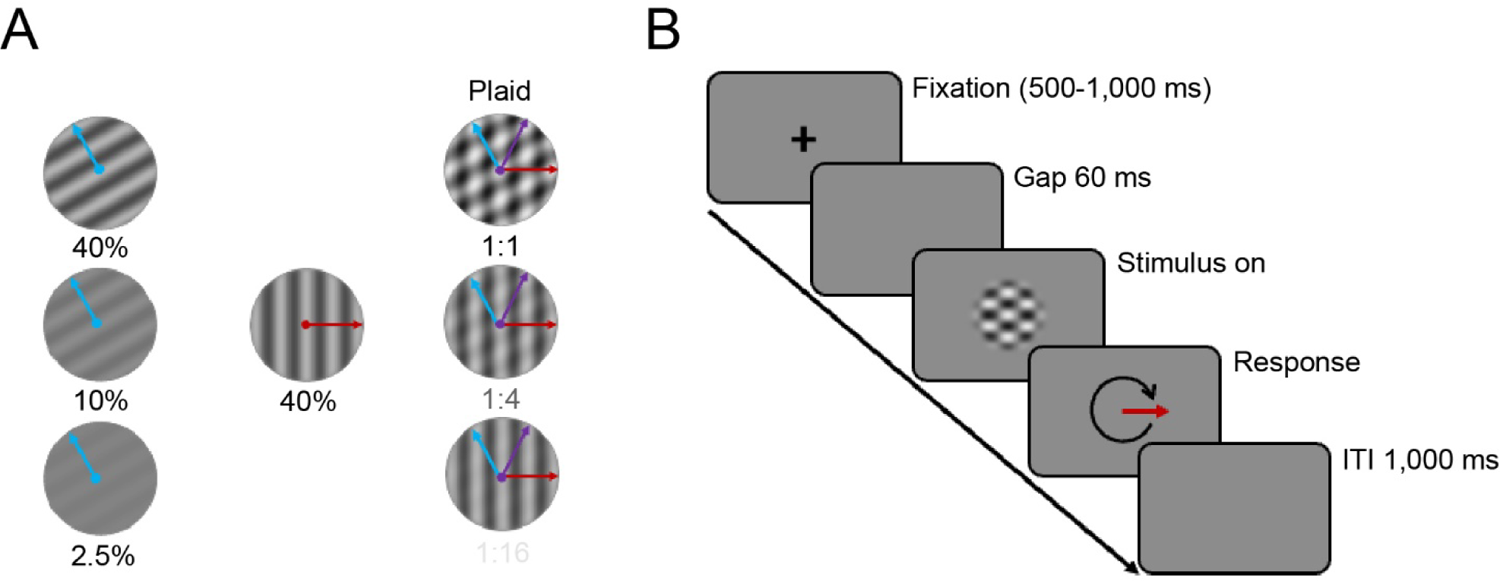
**(A)** Example plaids with a 1:1, 1:4, and 1:16 contrast ratio made by adding two gratings with different orientations and contrasts. Blue and red arrows indicate component motion directions; purple arrow shows pattern direction. **(B)** Trial timeline.

### Experimental design

At the beginning of each trial, observers fixated a centrally located cross for a randomly selected time between 500 and 1,000 ms. Following a 60 ms gap, we presented a drifting plaid or grating for a randomly-chosen duration of 70, 130, 250, or 500 ms. At the end of each trial, observers indicated the perceived stimulus direction by rotating an arrow on the screen using a trackball mouse. The next trial began after an inter-trial-interval of 1,000 ms (Fig. 1B). Randomly interleaved catch trials (1/7 of trials) contained no stimulus, and observers pressed the mouse button to continue to the next trial. We instructed observers to pay close attention to the stimulus, but they received no explicit eye movement instruction.

The computer selected plaid contrast ratio, presentation duration, and motion direction randomly for each trial. Each experimental session consisted of 420 trials, separated into 5 blocks of 84 trials each. Observers performed three one-hour sessions over three separate days. Across sessions, each observer completed a total of 1,260 trials, resulting in 45 repetitions per condition [45 × 7 (five plaids with different contrast ratios + high-contrast grating + catch trial) × 4 (presentation durations) = 1,260].

### Eye movement recording and data preprocessing

We analyzed eye movements within a time window of −50 ms to 500 ms from motion onset. We obtained eye velocity by differentiating eye position signals over time and smoothing using a second-order Butterworth filter (40 Hz cutoff frequency).

We removed saccades from eye movement traces for the OFR analysis. Saccades were defined as samples in which eye velocity exceeded a fixed velocity threshold of 30°/s for a minimum of 5 milliseconds; the nearest reversal in eye acceleration before eye velocity exceeded the threshold marked the saccade onset, and the nearest reversal after eye velocity returned below threshold marked the saccade offset.

For analysis and data presentation, we rotated the data so that the high contrast component motion was always at 0° (represented as horizontal). With the low-contrast component oriented 120° counter-clockwise from the high-contrast component, it follows that the rotated pattern motion direction (defined as the intersection of constraints; Adelson & Movshon, 1982) was always at 60° counter-clockwise relative to the high-contrast component (purple arrows in Fig. 1A). We refer to the 0° (high-contrast component) direction as ‘component’, the 90° direction as ‘orthogonal’, and the plaid direction (60°) as ‘pattern’.

We detected OFR onset by calculating least-squares fits of a 2-segment piecewise linear function to mean 2D eye velocity traces; the time of the break point defined the onset. We manually inspected all trials and excluded those with blinks, those in which the tracker lost the signal, or trials with undetected saccades within the first 300 ms of stimulus onset (3.1% of all trials). We also excluded trials in which observers reported a direction >60° from either the component or pattern direction (2.7% of all trials). One observer reported the opposite direction to the component or pattern motion in 16.4% of their trials. These trials were inconsistently spread across contrast ratios, presentation durations, and sessions; we excluded them from analysis. After applying the above exclusion criteria and eliminating catch-trials, 8,123 trials of 10,080 trials remained for analyses.

### Perceptual data and statistical analyses

We rotated observers’ perceptual judgements in the same way as their eye movements (i.e., treating 0° as the component direction, and 60° as the pattern motion direction), and collated responses for trials of each duration (70, 130, 250, and 500 ms). For the first three of these, we then compared perceptual responses to the direction of the OFR direction measured in 40 ms time-windows chosen to begin 25 ms after stimulus offset: 95-135 ms after stimulus onset (‘early OFR’), 155-195 ms (‘late OFR’), and 295-315 ms (‘tracking’) (Fig. 2A); we also collected behavioral reports after the longest trials, for which there was no corresponding OFR measurement. To quantify pattern sensitivity for OFR and perception, we calculated contrast ratio functions. We computed least-squares fits to the perceptual and eye movement data of the equation:

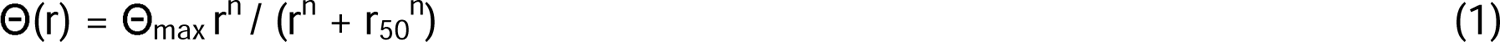

where Θ denotes the angle of the reported or eye movement direction, Θ_max_ the direction of the true motion of the stimulus (60°), r the contrast ratio, and r_50_ the contrast ratio at half asymptotic direction (30°). We used the r_50_ parameter to compare sensitivity to pattern motion direction between OFR and perception. Lower r_50_ values indicate higher sensitivity to pattern motion direction and higher values indicate a bias towards the high-contrast component.

**Figure 2.**
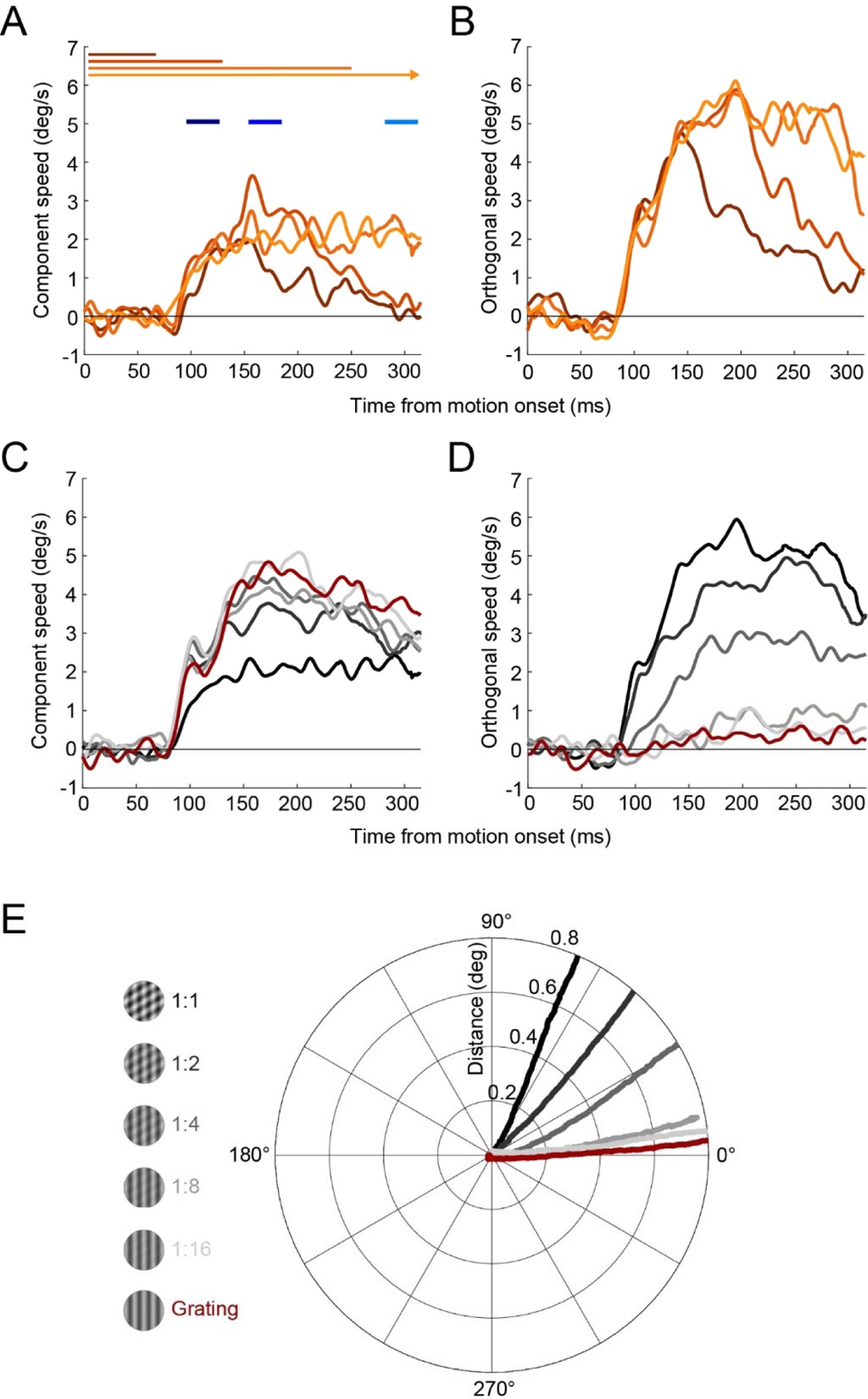
Single observer data, rotated as described in *Methods* so that eye movements in the direction of the high contrast component are labeled as component, and eye movements 90° away as orthogonal. **(A)** Component and **(B)** orthogonal speed over time, aligned to stimulus motion onset. Traces show averages across trials for each presentation duration. Blue lines indicate analysis time windows for early OFR (dark blue, 95-135 ms), late OFR (medium blue, 155-195 ms) and tracking (light blue, 275-315 ms). **(C)** Component and **(D)** orthogonal speed for each contrast condition (grey shades), for the two longest duration conditions. **(E)** Average eye movement trajectories for each contrast ratio.

We used single-trial estimates of reported and tracked direction to ask whether OFR and perception were correlated on a trial-by-trial basis. For this analysis, we applied an exclusion criterion so that only trials in which the eyes moved at a velocity of at least 1°/s and within 60° of either the component or pattern direction. We ran three linear mixed models (one per OFR analysis time window) with eye movement (early OFR, late OFR, or tracking), contrast ratio, their interaction, and observer as a grouping variable, allowing random intercepts and slopes for the correlation between eye movements and perception. All statistical analyses with an alpha level of 0.05, unless otherwise stated.

## Results

We recorded observers’ eye movements and motion direction estimates in response to drifting gratings or plaids of different contrast ratios. Figure 2 shows eye speed responses in the component (Fig. 2A) and orthogonal motion directions (Fig. 2B) for the observer with the strongest OFR (highest peak speed). This observer tracked stimuli at a latency of 83 ms after stimulus motion onset. For the shortest presentation duration (dark orange lines in Fig. 2A**,B**), OFR speed peaked at around 150 ms, and then started to decay. Speed peaked at ∼200 ms and then decayed for the 130-ms condition or was maintained for at least 320 ms for the two longest presentation durations (light orange lines). Based on these and similar traces for other observers, we chose to use data for all presentation durations for the early OFR measurements, data for the three longest durations for the late OFR measurements, and data for the two longest durations for the tracking measurements. Figures 2C and **2D** show component and orthogonal speed for the two longest presentation durations for the same individual observer. When presented with a single, high-contrast grating, this observers’ eyes moved solely in the component direction (zero orthogonal speed, Fig. 2D). With increasing contrast of the second component, component speed decreased and orthogonal speed increased, indicating a shift toward the pattern direction. Transforming 2D eye positions into polar coordinates (Fig. 2E) confirmed that the eyes moved in the 0°-direction in response to single gratings and for plaids with large ratios of component contrasts. The OFR was driven towards the pattern motion direction when component contrasts were within a factor of 2.

### Similar pattern motion sensitivity for OFR and perception

Eye movement responses were consistent across observers and closely resembled the results for the single observer shown in Figure 2. Across all observers, OFR was initiated with a mean latency of 80 ms (± 4 ms). Figure 3A shows relative eye direction over time across all observers. Before OFR onset (<80 ms), the eyes moved in random directions. Around the time of OFR onset, variability in eye movement direction decreased and the range of tracked directions collapsed to values between 0 and 60°, depending on contrast ratio. When presented with a single, high-contrast grating or a 1:16 plaid, the eyes moved in the direction of the high-contrast component. When both components had the same contrast (1:1 plaid), the eyes followed the pattern motion. For intermediate contrast ratios (1:8, 1:4, and 1:2), the eyes followed a weighted average of the two components. For these intermediate contrast ratios, the OFR direction shifted over time: during the early OFR (dark blue; Fig. 3A), eye movements were biased toward the high-contrast component—most pronounced for plaids with contrast ratios of 1:4 and 1:2 (darker shades of grey in Fig. 3A). During the late OFR and during tracking, eye movements were less biased towards the high-contrast component and more closely followed the pattern motion. This indicates a shift in the OFR from an early, component-driven response to a later, pattern-driven response that reflects a contrast-dependent weighted average of the directions of the two plaid components.

**Figure 3.**
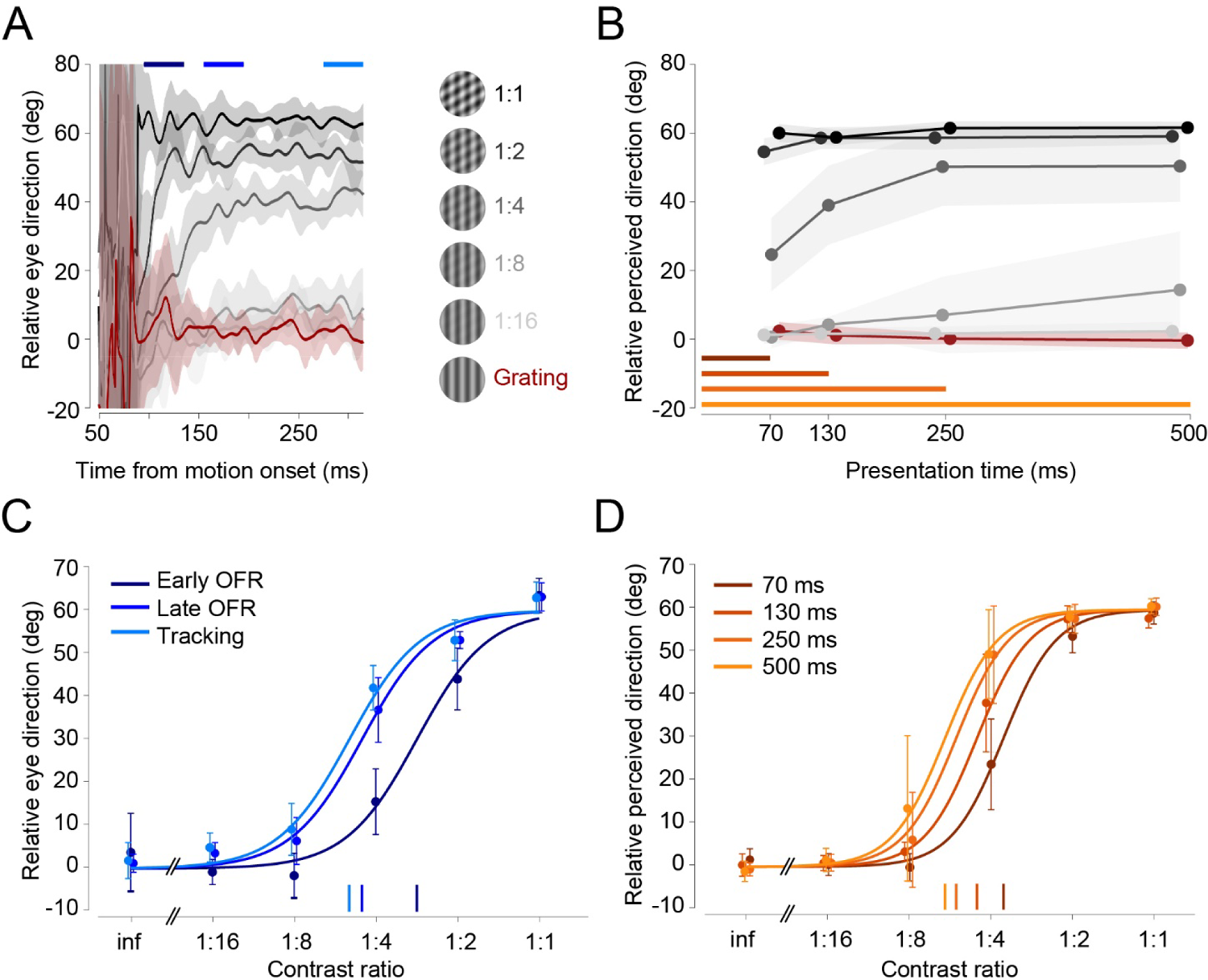
Comparison of contrast-dependent biases in OFR and perceptual estimates across observers (*n*=8). **(A)** Eye movement direction over time relative to stimulus onset for each contrast ratio. Traces show averages across observers and shaded areas represent ± 1 SD. **(B)** Reported motion direction for each contrast ratio and presentation duration.

Presenting stimuli for different durations allowed us to analyze perceptual reports as a function of time. Each data point in Figure 3B corresponds to one presentation duration. These results show that observers perceived component motion for the grating and the 1:16 plaid, and pattern motion for the 1:1 and 1:2 plaids. For intermediate contrast ratios, observers’ perception was biased by the high-contrast component when the stimulus was presented for 70 ms, but shifted toward pattern motion for longer presentation durations, in alignment with OFR results.

Shaded areas represent ±1 SD **(C)** Contrast ratio functions for early (dark blue), late (medium blue) and tracking (light blue) phases of the OFR. Dots and error bars show mean ± 1 SD across observers. Vertical lines on the baseline indicate r_50_. inf = grating. **(D)** Contrast ratio functions for perceived direction for each presentation duration. Same conventions as in panel C.

To directly compare pattern motion sensitivity between perceptual estimates and OFR, we plotted the tracked and reported direction as a function of contrast ratio and then fit contrast ratio functions. We extracted the contrast ratio at 50% of pattern direction (r_50_) as an indicator of pattern motion sensitivity. Figure 3C shows best fit contrast ratio functions for different phases of the OFR, using the means across all observers. A leftward shift of the function along the horizontal axis implies increasing pattern sensitivity from early to late OFR to tracking. We observed a similar increase in pattern sensitivity with increasing presentation duration for perceptual reports (Fig. 3D).

Figure 4 shows results of a condition-by-condition comparison of pattern motion sensitivity in perceptual reports vs. the OFR. A linear mixed model with observer as a grouping variable and random effects *intercept* and *slope* yielded a significant regression model (β = 0.674, *t*(30) = 6.70, *p* < .001), confirming a similar shift from early, component-driven to later, pattern-driven responses in eye movements and perception.

**Figure 4.**
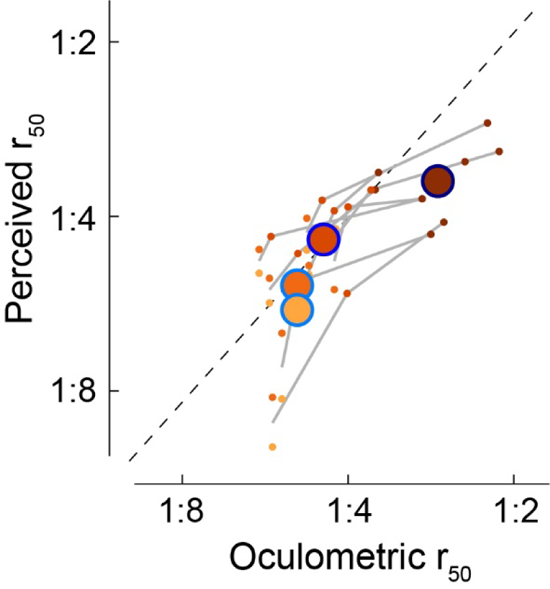
Comparison of biases in perception and tracking. Orange colors denote presentation durations, blue outlines are OFR phases. Grey lines connect individual observer data and show consistent time-shifted sensitivity across observers.

Comparing r_50_ values between the early phase of the OFR and perceptual reports in the shortest presentation duration condition revealed that observers’ earliest perceptual reports were more sensitive to pattern motion than the early OFR, indicated by lower r_50_ values for perceived direction (see dark red dots in Fig. 4). For all later analysis windows and longer presentation durations, pattern sensitivity was comparable between OFR and perceptual estimates (see yellow and orange dots in Fig. 4). Across all conditions, the exponents of the best fit contrast ratio functions were higher for perception (*M* = 5.92, *SD* = 1.54) compared to eye movements (*M* = 3.55, *SD* = 0.42; *t*(7) = 4.02, *p* = 0.005), indicating higher sensitivity in perception than in eye movements.

### Common noise sources for OFR and perception

We next analyzed the relationship between eye movements and perception on a trial-by-trial basis to determine whether variability in the OFR and in perceptual reports relies on common or private noise sources. Pooling data from all observers and trials for contrast ratios and OFR analysis intervals revealed larger variability in the OFR than in perceptual reports by a factor of 3. Within-subject variability was *M* = 37.4 (*SD* = 6.9; reported as across-subjects mean) for early OFR (Fig. 5, top row), *M* = 27.3 (*SD* = 5.1) for late OFR (middle row), and *M* = 28.8 (*SD* = 5.4) for tracking (bottom row). As Figure 5 reveals, variability was much smaller and largely constant across presentation durations for perceptual reports, ranging between *M* = 10.5 to 11.5.

**Figure 5.**
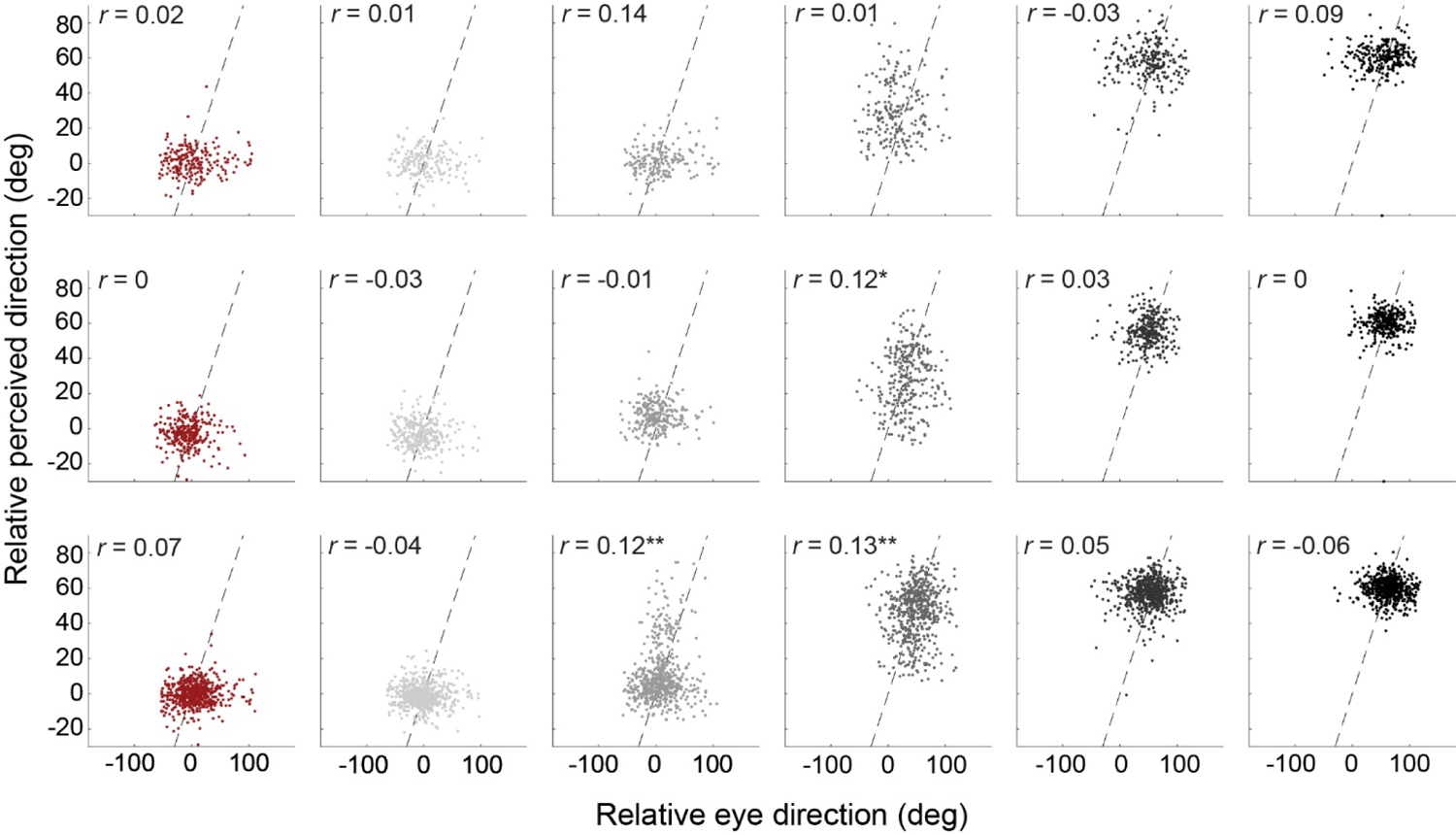
Trial-by-trial correlations between perceptual reports and OFR for all observers. Top row: early OFR (95-135 ms); middle row: late OFR (155-195 ms); bottom row: tracking (275-315 ms). Data for the tracking phase is pooled from the 250 and 500 ms presentation times. Dashed lines are identity lines.

In line with our observation that eye movements and perception showed similar pattern motion sensitivity during the later analysis window and longer presentation times, we observed trial-by-trial correlations between perceptual reports and OFR during the late tracking phase (Fig. 5, bottom row). Correlations were only observed for intermediate contrast ratios, likely because perceptual estimates were at ceiling with little to no variability in reported directions for low and high contrast ratios. These results were confirmed by linear mixed models for each analysis window, yielding a significant interaction between contrast and tracking only for the third analysis interval (*F*(5,3475.73) = 7.15, *p* < .001).

Whereas these results indicate that OFR and perceptual estimates might share the same sensory noise during later tracking and longer presentation times, we did not find significant correlations during the early OFR. A correlation during this early phase might be obscured by the large variability and low signal-to-noise ratio during early OFR.

## Discussion

We studied how the relative contrast of plaid components affects OFR and pattern motion perception and report three main results. First, plaids composed of gratings with equal luminance contrasts elicited an OFR in the direction of the pattern, aligned with observers’ perceived motion direction. For plaids with varying contrast ratios, we found a gradual shift in OFR direction tuning: early OFR was biased towards the high-contrast component whereas late OFR corrected the initial bias and moved the eyes towards a contrast-weighted average of the two plaid components. This indicates a shift in motion signal integration over time, from on early, component-driven response to a later, pattern-driven response (Pack et al., 2004; Smith et al., 2005). Second, perceived motion direction was biased by the high-contrast component when the stimulus was presented briefly, but shifted toward pattern motion for longer presentation durations to become aligned to the direction of late OFR and tracking. Third, for intermediate contrast ratios, eye tracking and perceptual estimates were correlated on a trial-by-trial basis.

### Shared pattern motion signals in eye movements and motion perception

Perception and tracking eye movements rely on motion processing in cortical visual areas MT and MST (Newsome et al., 1985; Newsome and Paré, 1988; Dürsteler and Wurtz, 1988; Kawano, 1999; Lisberger and Movshon, 1999; Takemura et al., 2002; 2007; Ilg and Thier, 2008). A subset of MT neurons signals the direction of pattern motion (Movshon et al., 1985; Rodman and Albright, 1989; Smith et al., 2005; Rust et al., 2006). These pattern-sensitive cells form the neural basis of coherent pattern perception (Adelson and Movshon, 1982; Movshon et al., 1985). When the components in a plaid have different luminance contrasts, human observers’ perception is biased towards the higher contrast component (Stone et al., 1990; Yo and Wilson, 1992; Bowns, 2013). A contrast-dependent perceptual bias is congruent with findings showing that sinusoidal grating contrast modulates perceived speed (Thompson, 1982) and the velocity gain of smooth pursuit eye movements (Spering et al., 2005). Contrast modulation of encoded component speed would result in contrast-dependent weighting when combining grating components to compute pattern motion direction (Stone et al., 1990). Indeed, the tuning curves of MT pattern cells in macaque monkeys are biased towards the higher-contrast component when responding to plaids composed of different contrast components (Kumbhani et al., 2008). Using the same plaid stimuli as Kumbhani and colleagues, we have now demonstrated a strong contrast-dependent bias in the earliest phase of the OFR, which gradually shifts towards the pattern motion direction for the late OFR and which matches perceptual reports.

Using plaids composed of one drifting and one stationary grating (type II or unikinetic plaids), Masson and Castet (2002) also observed an early component-driven response in the direction of the drifting grating and a subsequent pattern-driven response in the OFR. These behavioural observations are consistent with neurophysiological studies. Pack and Born (2001) measured the direction tuning of MT neurons to bar textures moving 45° obliquely to their orientation. Neural activity in area MT was initially dominated by component-motion (or ‘contour’) signals orthogonal to the bars. Over time, direction preference shifted and exhibited the true pattern (or ‘terminator’) motion direction (Lorenceau et al., 1993). Parallel findings and a rapid transition from component to pattern cell activity were observed with unikinetic plaids (Wallisch and Movshon, 2019). This change in direction tuning from component to pattern motion could be due to a combination of slower processing of low-contrast pattern (terminator) components (Majaj et al., 2002), and to a longer latency of pattern signals in MT (Pack et al., 2004; Smith et al., 2005). In our situation, a late-arriving pattern motion signal would help the oculomotor system to align the late phase of the OFR with pattern motion.

### Common and private sensory noise sources for OFR and perception

We observed trial-by-trial correlations between OFR and perception during the later tracking phase and for longer presentation times. Congruent with these findings, human smooth pursuit eye movements and perceptual direction judgments covary on a trial-by-trial basis (Stone & Krauzlis, 2003). Monkeys’ smooth pursuit variability during the initiation phase (first 140 ms of the response) is largely due to sensory noise that might be shared with perceptual responses (Osborne et al., 2005; 2007). Based on these findings, variability in eye movements and perception may result from a shared sensory noise source, with private noise sources specific to the perceptual and eye movement system added further downstream (Stone and Krauzlis, 2003; Lisberger, 2010). Figure 6 shows how this model may account for our results. Our data suggest that overall, OFR and perception rely on a similar readout of pattern motion signals. The computed signal is time-dependent and gradually shifts from a component-driven to a pattern-driven response (Pack and Born, 2001; Smith et al., 2005). Sensory noise may be shared by both effector systems and additional, effector-specific noise sources may act downstream. In line with our finding that trial-by-trial variability was larger for eye movements compared to perception, we propose that a private noise source underlies the variability in eye movements, which may include motor noise and larger measurement noise. We did not observe a correlation between OFR and perception for the early analysis window and the shortest presentation time. On the one hand, this may be the result of different motion processing for perception and early OFR (Simoncini et al., 2012), in line with our finding that perceptual reports were more sensitive to pattern motion than early OFR. On the other hand, the higher trial-by-trial variability during early OFR, compared to later OFR and tracking might have masked a correlation between eye movements and perception. During this early phase, eye speed is slow and prone to low signal-to-noise ratio.

**Figure 6.**
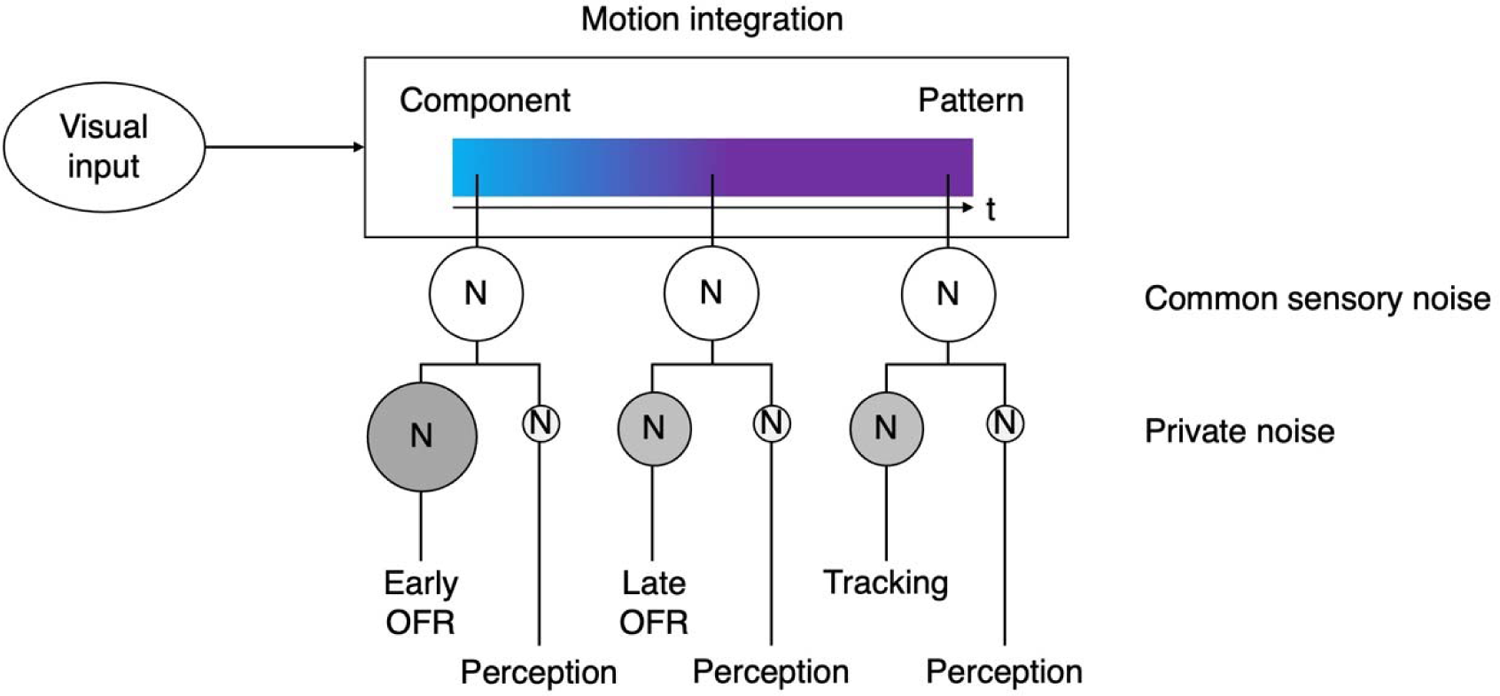
Schematic diagram summarizing key findings and proposed mechanisms. Plaids and gratings are processed via a motion integration stage providing a shared pattern motion signal for OFR and perception. Two noise sources are proposed: a common sensory noise source shared by perception and eye movements and a private source for each effector system. N = noise, size of the circles illustrates magnitude of intrasubject trial-by-trial variability.

## Conclusion

Our natural environment often contains complex motion patterns that must be captured and encoded by our visual and oculomotor systems. To understand how we operate in this challenging visuomotor world, it is critical to understand how pattern motion signals are computed to drive eye movements and perception. Here we show that ultra-short latency ocular following movements and perception integrate complex motion signals similarly, shifting from component-driven to pattern-driven responses during the first fraction of a second after the onset of visual motion. This also demonstrates that the OFR provides a sensitive and non-invasive tool to probe early visual motion processing and to study the time course of motion integration for complex signals.

## Acknowledgments

This research was funded by a Natural Sciences and Engineering Research Council of Canada (NSERC) Discovery Grant to MS (RGPIN 04987), by National Eye Institute Grants R01EY002017, R01EY004440, and P30EY13079 to JAM, and by a Robert Leet and Clara Guthrie Patterson Trust Postdoctoral Fellowship (to RDK)

## References

Adelson EH, Movshon JA (1982) Phenomenal coherence of moving visual patterns. Nature 300:523–525.

Bowns L (2013) An explanation of why component contrast affects perceived pattern motion. Vision Res 86:1–5.

Brainard DH (1997) The psychophysics toolbox. Spatial Vision 10: 433–436.

Cornelissen FW, Peters EM, Palmer J (2002) The Eyelink toolbox: eye tracking with MATLAB and the Psychophysics toolbox. Behav Res Method Instrum Comput 34(4):613–617.

Dürsteler MR, Wurtz RH (1988) Pursuit and optokinetic deficits following chemical lesions of cortical areas MT and MST. J Neurophysiol 60:940–965.

Gegenfurtner KR, Xing D, Scott BH, Hawken MJ (2003) A comparison of pursuit eye movements and perceptual performance in speed discrimination. J Vision 3: 865–876.

Gellman RS, Carl JR, Miles FA (1990) Short latency ocular-following responses in man. Vis Neurosci 5:107–122.

Glasser DM, Tadin D (2014) Modularity in the motion system: Independent oculomotor and perceptual processing of brief moving stimuli. J Vision 14(3):28,1–13.

Ilg UJ and Thier P (2008) The neural basis of smooth pursuit eye movements in the rhesus monkey brain. Brain Cogn 68:229–240.

Kawano K (1999) Ocular tracking: behavior and neurophysiology. Curr Opin Neurobiol 9:467–473.

Kumbhani RD, Saber GT, Majaj NJ, Tailby C, Movshon JA (2008) Contrast affects pattern direction selectivity in macaque MT neurons. Program No. 460.26. 2008 Neuroscience Meeting Planner. Washington, DC: Society for Neuroscience, 2008. Online.

Lisberger SG (2010) Visual guidance of smooth-pursuit eye movements: Sensation, action, and what happens in between. Neuron 65:477–491.

Lisberger SG, Movshon JA (1999) Visual motion analysis for pursuit eye movements in area MT of macaque monkeys. J Neurosci 19:2222–2246.

Lisi M, Cavanagh P (2015) Dissociation between the perceptual and saccadic localization of moving objects. Curr Biol 25:2535–2540.

Lorenceau J, Shiffrar M, Wells N, Castet E (1993) Different motion sensitive units are involved in recovering the direction of moving lines. Vision Res 33:1207–1217.

Majaj NJ, Smith MA, Kohn A, Bair W, Movshon JA (2002) A role for terminators in motion processing by macaque MT neurons? J Vision 2:415.

Masson GS, Castet E (2002) Parallel motion processing for the initiation of short-latency ocular following in humans. J Neurosci 22:5149–5163.

Masson GS, Perrinet LU (2012) The behavioral receptive field underlying motion integration for primate tracking eye movements. Neurosci Biobehav Rev 36:1–25.

Miles FA, Kawano K, Optican LM (1986) Short-latency ocular following responses of monkey. I. Dependence on temporospatial properties of visual input. J Neurophysiol 56:1321–1354.

Movshon JA, Adelson EH, Gizzi MS, Newsome WT (1985) The analysis of moving visual patterns. In: Pattern recognition mechanisms (Chagas C, Gattass R, Gross C, eds), pp 117–151. New York: Springer.

Movshon JA, Newsome WT (1996) Visual response properties of striate cortical neurons projecting to area MT in macaque monkeys. J Neurosci 16:7733–7741.

Newsome WT, Wurtz RH, Dürsteler MR, Mikami A (1985) Deficits in visual motion processing following ibotenic acid lesions of the middle temporal visual area of the macaque monkey. J Neurosci 3:825–840.

Newsome WT, Paré EB (1988) A selective impairment of motion perception following lesions of the middle temporal visual area (MT). J Neurosci 8:2201–2211.

Osborne LC, Hohl SS, Bialek W, Lisberger SG (2007) Time course of precision in smooth-pursuit eye movements of monkeys. J Neurosci 27(11):2987–2998.

Osborne LC, Lisberger SG, Bialek W (2005) A sensory source for motor variation. Nature 437:412–416.

Pack CC, Born RT (2001) Temporal dynamics of a neural solution to the aperture problem in visual area MT of macaque brain. Nature 409:1040–1042.

Pack CC, Gartland AJ, Born RT (2004) Integration of contour and terminator signals in visual area MT of alert macaque. J Neurosci 24:3268–3280.

Pattadkal JJ, Barr C, Priebe NJ (2023) Ocular following eye movements in marmosets follow complex motion trajectories. eNeuro 10(6): 1–9.

Pelli DG (1997) The VideoToolbox software for visual psychophysics: Transforming numbers into movies. Spatial Vision 10:437–442

Rodman HR, Albright TD (1989) Single-unit analysis of pattern-motion selective properties in the middle temporal visual area (MT). Exp Brain Res 75: 53–64.

Rust NC, Mante V, Simoncelli EP, Movshon JA (2006) How MT cells analyze the motion of visual patterns. Nat Neurosci 9:1421–1431.

Schütz AC, Braun DI, Gegenfurtner KR (2011) Eye movements and perception: a selective review. J Vis 11(5): 9.

Simoncini C, Perrinet LU, Montagnini A, Mamassian P, Masson GS (2012) More is not always better: adaptive gain control explains dissociation between perception and action. Nat Neurosci 15(11):1596–1603.

Smith MA, Majaj NJ, Movshon JA (2005) Dynamics of motion signaling by neurons in macaque area MT. Nat Neurosci 8(2):220–228.

Spering M, Carrasco M (2015) Acting without seeing: eye movements reveal visual processing without awareness. Trends Neurosci 38(4):247–258.

Spering M, Gegenfurtner KR (2007) Contrast and assimilation in motion perception and smooth pursuit eye movements. J Neurophysiol 97: 1353–1363.

Spering M, Montagnini A (2011) Do we track what we see? Evidence for common and independent processing of motion information for perception and smooth pursuit eye movements. Vision Res 51: 836–852.

Spering M, Kerzel D, Braun DI, Hawken MJ, Gegenfurtner KR (2005) Effect of contrast on smooth pursuit eye movements. J Vision 5(5):455–465.

Spering M, Pomplun M, Carrasco M (2011) Tracking without perceiving: a dissociation between eye movements and motion perception. Psychol Sci 22(2):216–225.

Stone LS, Watson AB, Mulligan JB (1990) Effect of contrast on the perceived direction of a moving plaid. Vision Res 30:1049–1067.

Stone LS, Krauzlis RJ (2003) Shared motion signals for human perceptual decisions and oculomotor actions. J Vision 3: 725–736.

Takemura A, Inoue Y, and Kawano K (2002) Visually driven eye movements elicited at ultra-short latency are severely impaired by MST neurons. Ann NY Acad Sci 956:456– 459.

Takemura A, Murata Y, Kawano K, Miles FA (2007) Deficits in short-latency tracking eye movements after chemical lesions in monkey cortical areas MT and MST. J Neurosci 27:529–541.

Tavassoli A, Ringach DL (2010) When your eyes see more than you do. Curr Biol 20:R93– R94.

Thompson P (1982) Perceived rate of movement depends on contrast. Vision Res 22:377–380.

Wallisch P, Movshon JA (2019) Responses of neurons in macaque MT to unikinetic plaids. J Neurophysiol 122:1937–1945.

Yip HMK, Allison-Walker TJ, Cloherty SL, Hagan MA, Price NSC (2023) Ocular following responses of the marmoset monkey are dependent on postsaccadic delay, spatiotemporal frequency, and saccade direction. J Neurophysiol 130:189–198.

Yo C, Wilson HR (1992) Perceived direction of moving two-dimensional patterns depends on duration, contrast and eccentricity. Vision Res 32:135–147.

